# Differential activation of mouse and human Panx1 channel variants

**DOI:** 10.1101/2023.10.11.561880

**Authors:** Antonio Cibelli, Preeti Dohare, David C. Spray, Eliana Scemes

## Abstract

Pannexins are ubiquitously expressed in human and mouse tissues. Pannexin 1 (Panx1), the most thoroughly characterized isoform of this family, forms plasmalemmal membrane channels permeable to relatively large molecules, such as ATP. Although human and mouse Panx1 amino acid sequences are conserved in the presently known regulatory sites involved in trafficking and modulation of the channel, differences are reported in the N- and C-termini of the protein, and the mechanisms of channel activation by different stimuli remain controversial. Here we used a neuroblastoma cell line to study the activation properties of endogenous mPanx1 and exogenously expressed hPanx1. Dye uptake and electrophysiological recordings revealed that in contrast to mouse Panx1, the human ortholog is insensitive to stimulation with high extracellular [K^+^] but responds similarly to activation of the purinergic P2X7 receptor. The two most frequent Panx1 polymorphisms found in the human population, Q5H (rs1138800) and E390D (rs74549886), exogenously expressed in Panx1-null N2a cells revealed that regarding P2X7 receptor mediated Panx1 activation, the Q5H mutant is a gain of function whereas the E390D mutant is a loss of function variant. Collectively, we demonstrate differences in the activation between human and mouse Panx1 orthologs and suggest that these differences may have translational implications for studies where Panx1 has been shown to have significant impact.

## INTRODUCTION

The Pannexin (Panx) family of proteins was discovered in the early 2000s, through *in silico* search for mammalian orthologs of the invertebrate gap junction forming proteins, the innexins (to which pannexins have <20% identity) [1]. Three members belong to Panx family (Panx1, Panx2 and Panx3). Panx1 is the most studied, forming large conductance channel allowing the passage of molecules up to 1 KDa, including fluorescent dyes and ATP [2, 3]. Each subunit of the channel pore consists of four transmembrane domains, cytosolic N- and C-termini, a single intracellular loop and two extracellular loops with a site for N-glycosylation [1, 4–6]. Although many innexins form intercellular channels, it has been speculated that the Panx1 N-glycosylation may prevent the pannexons from adjacent cells to do so, although whether this incompatibility is absolute still is a matter of debate (reviewed in [7]).

Panx1 is ubiquitously expressed in human and mouse tissues [4] and in both neurons and glia in the nervous system [8–13]. Multiple mechanisms have been reported to activate Panx1 channels [14], and stimuli that promote Panx1 channel activity include membrane stretch and hypotonicity [3, 15, 16], increased intracellular calcium levels [15], receptor mediated pathways (NMDA, P2X7, alpha-adrenergic, insulin, CXCR4: [2, 17–23]), and caspase3- mediated C-terminal cleavage [24]. However, there are still open questions regarding whether high extracellular [K^+^] affects Panx1 channel gating (see [14, 25]). Part of the reason for this uncertainty is that studies by different laboratories have focused on either rodent or human Panx1 orthologs, where amino acid differences can add another layer of complexity when analyzing gating mechanisms. Interestingly, in this regard is the finding that the most frequent missense *PANX1* variant in the human population (Q5H; rs1138800, where glutamine (Q) is substituted with Histidine (H) at amino acid position 5) is associated with a gain-of-function of the channel [26]. Another, very rare, polymorphism (R217H, rs143240087, in which an Arginine (R) is substituted with Histidine (H) at position 217) is associated with multisystem dysfunction [27] and encodes a nonfunctional channel.

To evaluate the extent to which mouse and human Panx1 channel activation modes may differ from each other and to evaluate potential differences between hPanx1 channel variants, we performed electrophysiological and dye uptake assays on cells exogenously expressing these transcripts. Here we show that in contrast to mouse, human Panx1 is insensitive to moderate elevation in extracellular [K^+^] but responds similarly to P2X7 stimulation. Characterization of the two most frequent Panx1 channel polymorphisms found in the human population revealed that, regarding P2X7 receptor mediated Panx1 activation, the Q5H isoform is a gain of function variant and that the E390D isoform is a loss of function variant. The differences reported here between the modes of activation between Panx1 orthologs are likely to have important implications to translational studies into disorders such as epilepsy and chronic pain, where Panx1 activation has been shown to have significant impact.

## MATERIALS AND METHODS

### Cells

As previously described [28], the mouse neuroblastoma cell line (ATCC CCL-131; here termed N2a) was cultured in Dulbecco’s modified essential medium (DMEM) supplemented with 10% fetal bovine serum and 5% penicillin/streptomycin. Deletion of endogenous mPanx1 was achieved by CRISPR editing [28] and Panx1-null cell lines stably expressing human Panx1 variants were generated using lentiviral constructs, as described below.

### Generation of hPanx1 N2a clones

Recombinant lentivirus particles pLV[Exp]- mcherry:T2A-Puro-EFS (VectorBuilder, Chicago, IL, USA) were used to express hPanx1 (NM_015368.3) and two of its variants, hPanx1Q5H (rs1138800) and hPanx1E390D (rs74549886). Cells were infected using three multiplicities of infections (MOI: 2.5, 5.0 and 10.0) for each of the vectors and cells maintained for 24 hrs at 37°C in a humidified 5% CO2 incubator. After removal of medium containing the virus, cells were maintained in regular DMEM for 24 hrs and then maintained in DMEM containing puromycin for selection of clones expressing the mCherry marker, visualized using an epifluorescence microscope.

### Quantitative Polymerase Chain Reaction (qPCR)

The qPCR was performed using SYBR GREEN Master Mix (Applied Biosystems) in an Applied Biosystems 7300 Real-Time PCR System (Forester City, CA), according to the manufacturer’s instructions. Briefly, 2 μg total RNA was reverse transcribed into cDNA using SuperScript VILO cDNA Synthesis (Invitrogen). Amplification was carried out for no more than 28 cycles with annealing temperature of 60°C in 25 μl final volume. Finally, a dissociation profile of the PCR product(s) was obtained by a temperature gradient running from 60°C to 95°C. The primers used for qPCR are shown below. Two technical replicas for each experiment with total of 4 biological replicas for each N2a clone were amplified. Relative gene expression levels of the mRNA analyzed were calculated using the ΔΔCT method, where values obtained for the gene of interest were first normalized to those of the reference, housekeeping gene, GAPDH, and subsequently to those of their respective controls (mouse brain and the human HeLa cells). Validation of the comparative Ct method was obtained after estimation of equality of efficiencies of target and reference amplifications (ΔCt (Cttarget – Ctreference)) over serial sample dilutions.

*Human PANX1 primers*:

F: TTTACAACCGTGCAATTAAGGCT R: AAGTTCTCGGTAACACCTGGA

*Mouse Panx1 primers*:

F: caggctgcctttgtggattc R: cgggcaggtacaggagtatg

### Immunocytochemistry

N2a cells plated on glass bottomed dishes were immunostained using the chicken anti-Panx1 (1:500; extracellular loop (CZVQQKSSLQSES); Aves Lab #6358, Davis, CA, USA). Cells fixed in 4% p-formaldehyde (PFA) were incubated for 30 min in a blocking solution (10% goat serum in PBS) and overnight with chicken anti-Panx1 antibody diluted (1:500) in blocking solution. After washout of primary antibody, cells were exposed to goat anti-chicken secondary antibody (1:2000; Alexa Fluor 488; Invitrogen, Waltham, MA, USA) for 1 hr at room temperature, prior to mounting (Vectashield containing DAPI; Vector Laboratory, Newark, NJ, USA). Images were captured with MetaFluor software (Molecular Devices, CA, USA) using a CoolSNAP-HQ2 CCD camera (Photometrics, AZ, USA) attached to an inverted Nikon microscope (Eclipse TE-2000E, Nikon, Japan) equipped with a 20X dry objective and FITC filter set.

### Electrophysiology

Cells were plated on glass coverslips 24 hrs prior to recordings. Whole cell patch clamp recordings were performed on cells that were round and exhibited no or few short processes, as previously described [29]. Briefly, cells were bathed in Krebs solution containing (mM): NaCl 147, HEPES 10, glucose 13, CaCl2 2, MgCl2 1 and KCl 2.5, pH 7.4; [K^+^] was varied through equimolar substitution of NaCl with KCl. The pipette solution contained (mM): CsCl 130, EGTA 10, HEPES 10, CaCl2 0.5, pH 7.4. Activation of Panx1 channels by voltage was performed by applying 5.5 sec. voltage ramps from a holding potential of −60 mV to +80 mV. To analyze the participation of Panx1 channels in agonist-induced P2X7R activation, N2a cells were bathed in zero Ca^2+^/Mg^2+^ phosphate buffered solution (PBS) supplemented with 5.5 mM glucose and the P2X7R agonist BzATP (100 μM; Sigma-Aldrich, MO, USA) was superfused for 5–10 sec. Panx1 currents induced by BzATP were recorded while applying the same voltage ramp protocol described above. Electrophysiological recordings were accomplished using an Axopatch 200B amplifier and pClamp10 software (San Jose, CA, USA) was used for data acquisition and analysis. Changes in peak conductance induced by 10 mM K^+^ and by BzATP were normalized to those recorded in control conditions and expressed as fold changes.

### Dye uptake

Opening of Panx1 channels was evaluated by the dye uptake method, as previously described [8, 9, 30]. Briefly, N2a cells plated on glass bottomed dishes were incubated for 10 min in zero Ca^2+^/Mg^2+^ PBS containing 5 µm YoPro (YOPRO-1 iodide; Thermo-Fisher, USA) in the absence and presence of 100 µM BzATP. For some experiments, cells were treated with the Panx1 channel blocker, mefloquine (MFQ, 10 µM; Bioblock, San Diego, CA, USA) prepared in zero Ca^2+^/Mg^2+^ PBS containing 5 µm YoPro, with and without BzATP. To evaluate the extent to which high K^+^ solution led to Panx1 channel opening, cells were exposed for 10 min to 2.5 mM or 10 mM K^+^ Krebs solution containing 5 µM YoPro. After removal of the dye containing solution, cells were fixed in 4% PFA for 5 min and then YoPro fluorescence intensity measured from images captured using Eclipse TE-2000E Nikon (Nikon, Japan) or an BZ-X800 Keyence (Keyence Corporation of America, Itasca, IL, USA) inverted microscopes equipped with 20x objective and FITC filter sets. Fluorescence intensity was measured from areas of interest (ROI) placed on cells using MetaFluor software or Fiji-ImageJ. The influx of YoPro in BzATP-treated and high K^+^ exposed cells was calculated as the dye intensity value relative to that recorded under respective control conditions (BzATP compared to untreated or 10 mM compared to 2.5 mM K^+^).

### Statistical analyses

GraphPad Prism-8 software (San Diego, CA, USA) was used for statistical analyses. One-way ANOVA followed by Sidak’s or Tukey’s multiple comparison tests was used to determine significant difference between means. In some cases, t-test were used to analyze differences between 2 groups. Values are expressed and mean ± SE, with p<0.05 considered as statistically significant.

## RESULTS

### 1. The mouse neuroblastoma cell line (N2a) expresses endogenous Panx1

The mouse neuroblastoma cell line, N2a, expresses endogenous Panx1 as evidenced by q-PCR and immunocytochemistry (Figures 1A, B). Compared to the mouse brain, N2a cells express substantially lower Panx1 transcript levels (Figure 1A). Immunocytochemistry performed in non-permeabilized N2a cells using a Panx1 antibody to the extracellular loop revealed that Panx1 is present in this cell line (Figure 1B).

**Figure 1.**
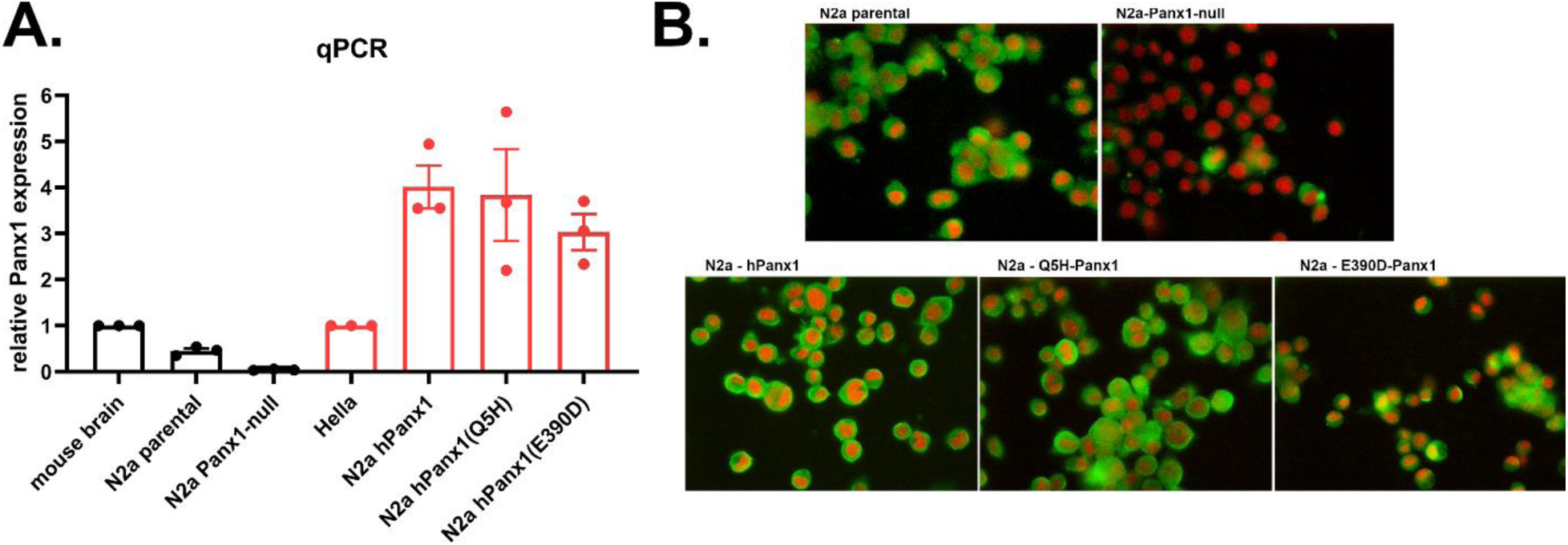
Expression levels of mouse and human Panx1. (**A**) Mean ± SEM values of the relative expression levels of Panx1 transcripts obtained by q-RT-PCR performed in N2a cell expressing endogenous mPanx1 (parental), N2a lacking Panx1 (Panx1-null), and in N2a expressing human wildtype Panx1 (hPanx1) and two variants (Q5H, E390D). Mouse and human Panx1 mRNA levels were normalized to those obtained from mouse brain and HeLa cells, respectively. (**B**) Representative immunofluorescence images obtained from N2a parental, Panx1-null and Panx1- null N2a cells expressing human (h)Panx1 constructs. Immunocytochemistry was performed on non-permeabilized cells using an extracellular epitope anti-Panx1 antibody (shown in green) and cells counterstained with DAPI (in red). Scale bar: 20 µm.

For studies described below using N2a exogenously expressing hPanx1, we first generated a N2a-Panx1-null cell line using Crispr/Cas-9 technology. Using this approach, we obtained successful knockdown of Panx1, as indicated by almost complete absence of Panx1 transcript as measured by q-PCR (Figure 1A). Immunocytochemistry revealed complete knockdown of Panx1 in the vast majority of the N2a cells although a few still displayed Panx1 protein (Figure 1B).

### 2. Mouse Panx1 channels are sensitive to extracellular [K^+^] and to P2X7 receptor stimulation

Whole cell voltage-clamp recordings performed on cultured N2a cells bathed in 2.5 mM K^+^ solution revealed the presence of outwardly rectifying currents at depolarizing potential (Figure 2A, gray line). These currents are similar to those previously recorded from N2a cells transfected with mPanx1 [29] and to those from cultured mouse astrocytes [9]. These endogenous outward currents recorded from N2a cells which we have previously shown to be sensitive to Panx1 channel blocker mefloquine (MFQ; [29]) were substantially lower in Panx1-null N2a cells (Figure 2A); the mean peak conductance (1.66 ± 0.17 nS, n = 4 cells) of Panx1-null cells measured at +80 mV was 2.7 fold smaller than that recorded from N2a expressing endogenous Panx1 channels (4.51 ± 0.44 nS, n = 7 cells). To test whether endogenous Panx1 current in N2a cells was also sensitive to elevated extracellular K^+^ and activated following stimulation of the P2X7 receptor as we have previously reported in N2a cells over-expressing mPanx1 [31], electrophysiological and dye uptake assays were performed. Voltage-activated outwardly rectifying Panx1 currents recorded from N2a cells expressing endogenous Panx1 were potentiated by the superfusion of solution containing 10 mM K^+^ (equimolar Na^+^ replacement) and were decreased in Panx1-null cells (Figure 2A). In N2a parental cells, bath application of 10 mM K^+^ induced a 1.85 ± 0.15 (n = 5 cells) fold increase in peak conductance compared to that measured when these cells were bathed in 2.5 mM K^+^ solution (Figure 2B); in contrast, compared to 2.5 mM K^+^ solution, the peak conductance recorded from Panx1- null cells decreased slightly to 0.77 ± 0.04 fold (n = 4 cells) when cells were exposed to 10 mM K^+^ solution (Figure 2B). Similarly, bath application of 100 µM BzATP to N2a parental cells increased peak conductance by 2.03 ± 0.22 fold (n = 5 cells) and in Panx1- null cells this treatment minimally affected peak conductance decreasing it to 0.86 ± 0.10 fold (n = 5 cells) compared to that recorded in the absence of the agonist (Figures. 2C, D).

**Figure 2.**
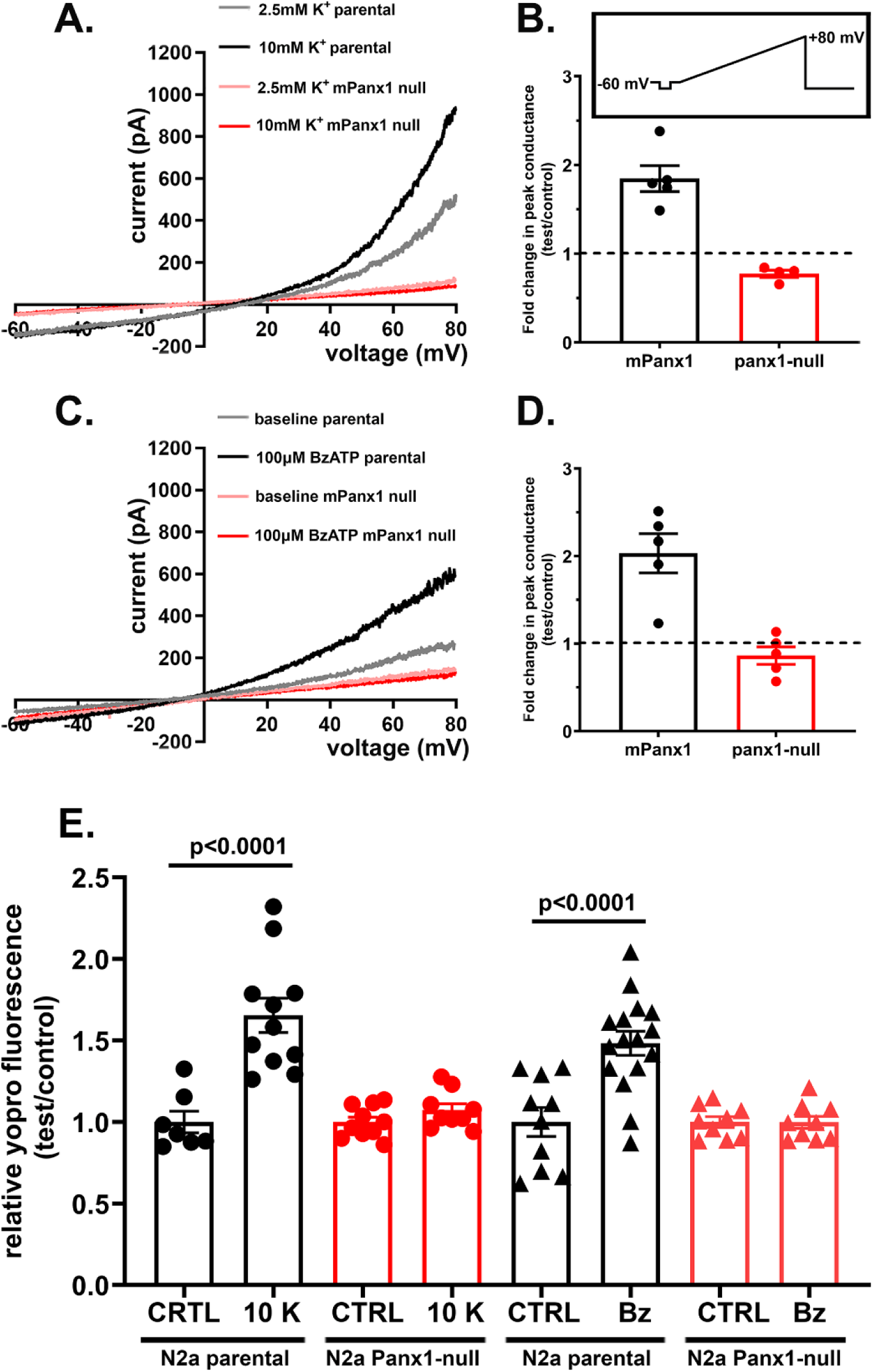
Mouse Panx1 conductance is activated by elevated extracellular [K^+^] and following P2X7R stimulation. (**A**-**D**) Electrophysiological recordings and summary histograms obtained from parental N2a and Panx1-null N2a cells exposed to (**A**-**B**) 10 mM K^+^ and (**C**-**D**) 100 µM BzATP. Note in parts A and C that N2a parental cells responded to high K^+^ solution and to BzATP with increased current amplitudes (gray and black curves) in response to 5.5 sec long voltage ramps from -60 to +80 mV (inset in panel **B**), while no changes in current amplitudes were recorded from Panx1-null N2a cells (pink and red curves, which totally overlap). The mean ±SEM values of the fold changes in peak conductance induced by 10 mM K^+^ and BzATP obtained for parental and Panx1-null N2a cells are shown in parts **B** and **D**, respectively. N= 4 - 5 cells. (**E**) Mean ± SEM values of the relative YoPro fluorescence intensity obtained from parental and Panx1-null N2a cells after 10 min exposure to 2.5 or 10 mM K^+^ solutions and to 100 µM BzATP. Values were normalized to those obtained under control conditions. Each point on the graphs represents mean value of relative YoPro fluorescence changes recorded from all cells present in each field of view obtained from at 3 – 5 independent experiments. p values obtained from ANOVA followed by Sidak’s multiple comparison test.

Parallel dye uptake experiments indicated that after 10 min exposure to 10 mM K^+^ solution or to 100 µM BzATP YoPro fluorescence intensity increased significantly in parental N2a cells but not in Panx1-null N2a cells (Figure 2E; Supplementary Figure S1). Compared to that measured under control conditions, high K^+^ solution or bath application of 100 µM BzATP increased YoPro fluorescence intensity in N2a parental cells by 1.66 ±0.11 fold (n = 11 fields) and 1.48 ± 0.07 fold (n = 16 fields), respectively, (Figure 2E). In Panx1-null cells, no changes in YoPro fluorescence intensity were recorded 10 min after exposure to high K^+^ solution (1.07 ± 0.04 fold, n = 10 fields) or to bath application of 100 µM BzATP (1.00 ± 0.04 fold, n = 9 fields; Figure 2E; Supplementary Figure S1).

Thus, these results clearly show that in N2a cells, endogenous mPanx1 channels are sensitive to extracellular [K^+^] and can be activated following stimulation of the P2X7 receptor. Moreover, our electrophysiological and dye uptake results indicate that increases in Panx1 mediated ionic currents caused by these agents correspond to increased influx of the moderately large cationic fluorescent dye YoPro.

### 3. Properties of human Panx1 channels differ from mouse Panx1

Comparison between human and mouse Panx1 amino acid sequences reveal differences between the species, particularly at the second half of the protein (15 differences in N-terminal 210 amino acid portion *vs* 40 differences in the C-terminal 210 amino acid portion; Figure 3A). However, the presently known regulatory sites in the intracellular loop and the C-terminal domains involved in Panx1 trafficking and channels modulation are conserved [32, 33]. This implies that mouse and human Panx1 would be likely to respond similarly to gating stimuli. Nevertheless, reports have indicated that differently from mouse, hPanx1 channels are constitutively closed, only becoming activated following caspase cleavage at position 371 of the C-terminus [24]. In addition, missense variants in PANX1 have been associated with human diseases. These variants include rs1138800 (Q5H) [26, 34, 35] which leads to a hyperfunction of the channels and rs143240087 (R217H) which was reported to have a dominant-negative effect [27].

**Figure 3.**
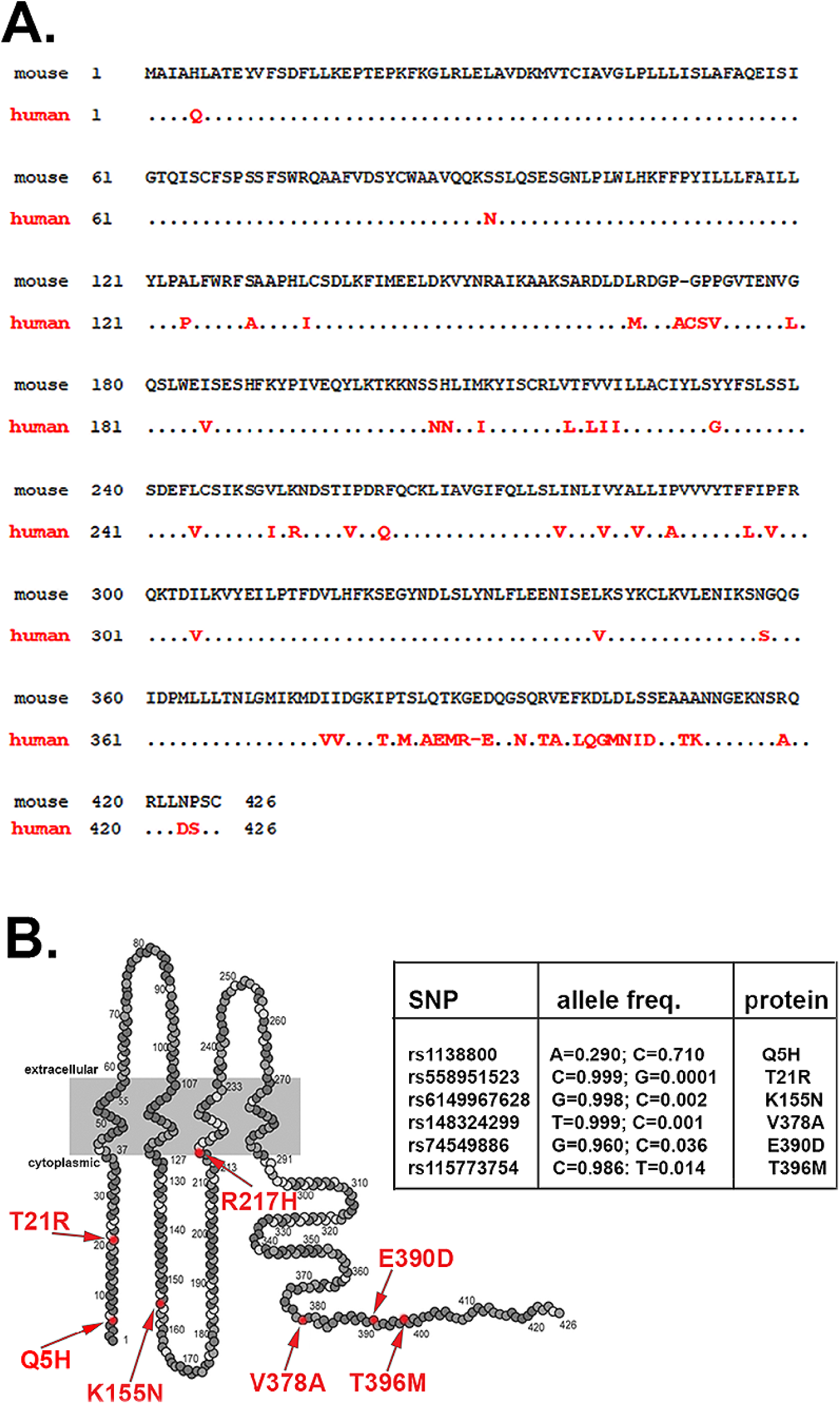
Sequences of mouse and wildtype human Panx1 and variants. (**A**) Comparison between amino acid sequences of mouse and human pannexin proteins. In red are amino acids found in human Panx1 that differ from those found in mouse. (**B**). Human Panx1 topology showing most frequent missense variants reported (arrows) and table showing the SNP allele frequencies.

Inspection of the polymorphism data base (dbSNPs: https://www.ncbi.nlm.nih.gov/snp/) reveals that among the several missense PANX1 variants located in the cytoplasmic domain, two are more frequently found in the human population than others, Q5H (rs1138800) and E390D (rs74549886) (Figure 3B). According to the TOPMed project the Q5H frequency of the reference and altered alleles A>C are 0.28 and 0.72, respectively; while the E390D frequency of the reference and altered allele G>T are 0.96 and 0.038, respectively.

To evaluate whether activation modes of hPanx1 and two of its variants differed from mPanx1, we used Panx1-null N2a cells infected with lentivirus constructs to express wild-type hPanx1, and the Q5H and the E390D variants. Of the three multiplicities infections used (MOI), MOI of 10 resulted in consistent and homogenous expression of hPanx1 and its variants. The mRNA and protein expression levels of each of the human Panx1 variants in Panx1-null N2a cells were similar (Figures 1A, B) and mRNA levels were higher than that of the endogenous transcript in the human HeLa cell line (Figure1A).

Electrophysiological recordings indicated that infection of N2a cells with all three hPanx1 constructs in lentivirus vectors led to outwardly rectifying currents when bathed in 2.5 mM K+ solution (Figure 4); mean peak conductance of hPanx1 (2.53 ± 0.53 nS, n = 6 cells), Q5H (2.08 ± 0.19 nS, n = 6 cells), and E390D (2.84 ± 0.17 nS, n = 5 cells) were not statistically different (p=0.39, ANOVA). However, the mean conductance of hPanx1 channel and that of its variants were significantly smaller than those measured from mPanx1 (4.51 ± 0.44 nS, n = 7 cells; p=0.001, ANOVA). Also, differently from mPanx1, hPanx1 and its variants were insensitive to 10 mM K^+^ (compare Figures 4A-D to Figures 2A-D). No significant changes in peak conductance were recorded when N2a cells expressing hPanx1 (0.98 ± 0.06; n = 6 cells), Q5H-Panx1 (0.92 ± 0.04; n = 9 cells), and E390D-Panx1 (0.91 ± 0.07; n = 5 cells) were exposed to 10 mM K^+^ solution (Figure 4D).

**FIGURE 4.**
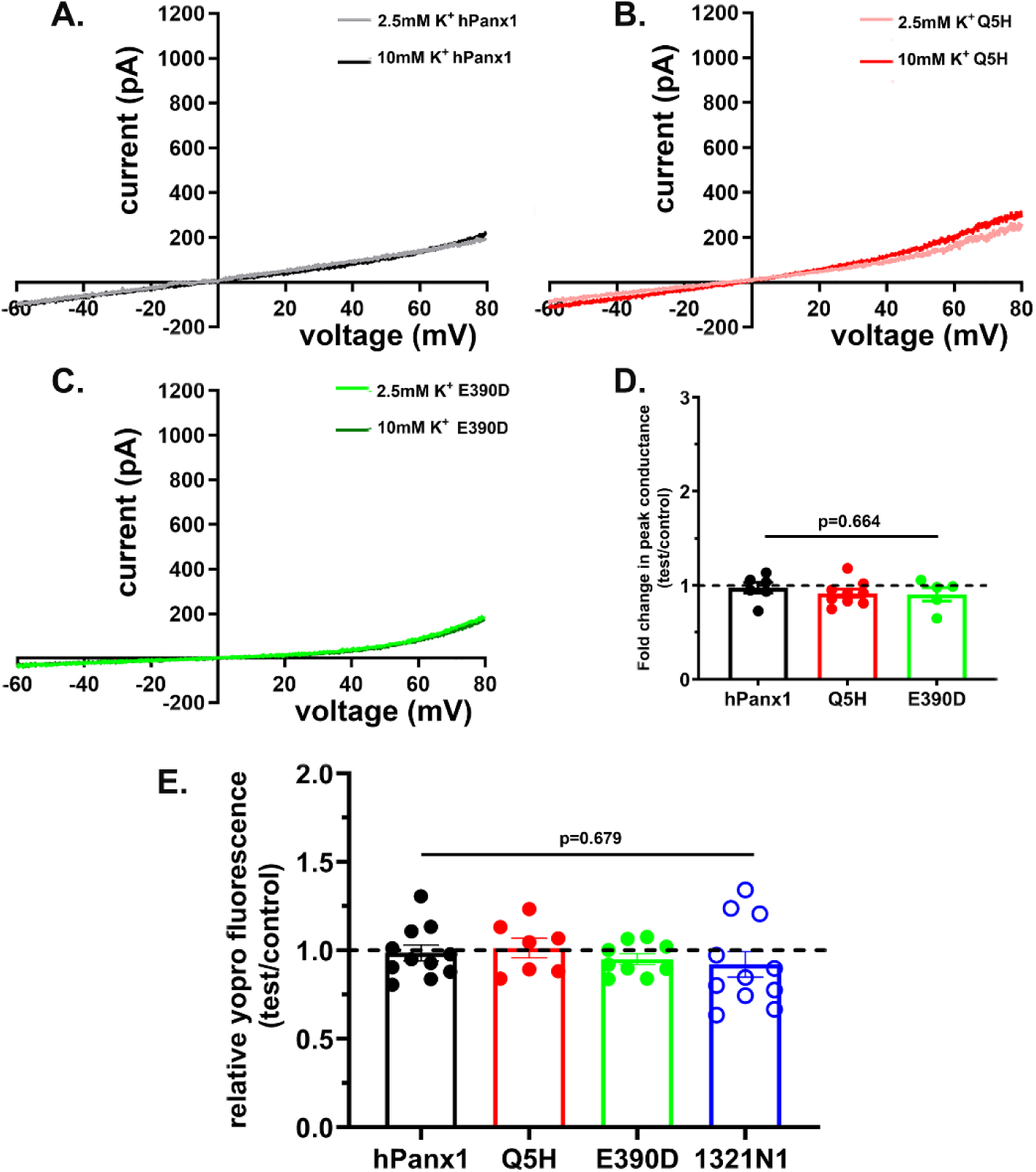
Human Panx1 channels are insensitive to elevated extracellular [K^+^]. (**A**-**C**) Examples of Panx1 currents recorded from N2a cells expressing hPanx1 and two of its variants (Q5H and E390D) exposed to solutions containing 2.5 and 10 mM K^+^. Note that increased extracellular K^+^ concentration did not lead to increase in Panx1 currents in any of these sequence variants. (**D**) Mean ± SEM values of fold changes in peak conductance obtained from hPanx1, Q5H and E390D Panx1 channels induced by 10 mM K^+^ solution. N = 4 - 9 cells. (**E**) Mean ± SEM values of the relative YoPro fluorescence intensity changes obtained from N2a cells expressing hPanx1 and two variants (Q5H and E390D), and from the human astrocytoma cell line 1321N1 that express endogenous hPanx1 exposed for 10 min to 10 mM K^+^. Dashed line represents mean values of fluorescence recorded from cells exposed to 2.5 mM K^+^ solution. Each point on the graphs represents a mean value of YoPro fluorescence recorded from all cells present in a field of view obtained from at 3 – 5 independent experiments.

As expected from electrophysiological recordings, dye uptake assay showed that exposure to 10 mM K^+^ solution did not lead to influx of YoPro dye in N2a cells expressing hPanx1 (Supplementary Figure S1) or its variants (Figure 4E). After 10 min exposure to high K^+^, the relative YoPro fluorescence intensity change recorded from hPanx1 was 0.99 ± 0.04 (n = 11 fields), from Q5H was 1.01 ± 0.06 (n = 7 fields) and from E390D was 0.95 ± 0.03 (n = 9 fields). To evaluate whether this K^+^-insensitivity might be due to post-translational modifications that could occur when human proteins are expressed in a mouse cell line, we performed similar dye uptake experiments using the human astrocytoma cell line 1321N1, which expresses endogenous hPanx1 (Silverman et al., 2009). As recorded from N2a expressing hPanx1, the 1321N1 cells were also insensitive to 10 mM K^+^ (0.92 ± 0.08, n = 11 fields; Figure 4E).

We then evaluated whether human Panx1 responded to P2X7 receptor stimulation. Electrophysiological recordings revealed that in contrast to their K^+^- insensitivity, hPanx1 channels were activated following stimulation of the P2X7 receptor with BzATP (Figures 5A-D). Exposure to 100 µM BzATP induced a 2.05 ± 0.24 fold change in peak conductance in N2a cells expressing hPanx1 and 3.49 ± 0.29 fold change in N2a expressing Q5H-Panx1 (Figure 5D). Interestingly, the E390D variant did not respond to P2X7R activation (change in peak conductance: 1.02 ± 0.07 fold, n = 8 cells; Figure 5D).

**FIGURE 5.**
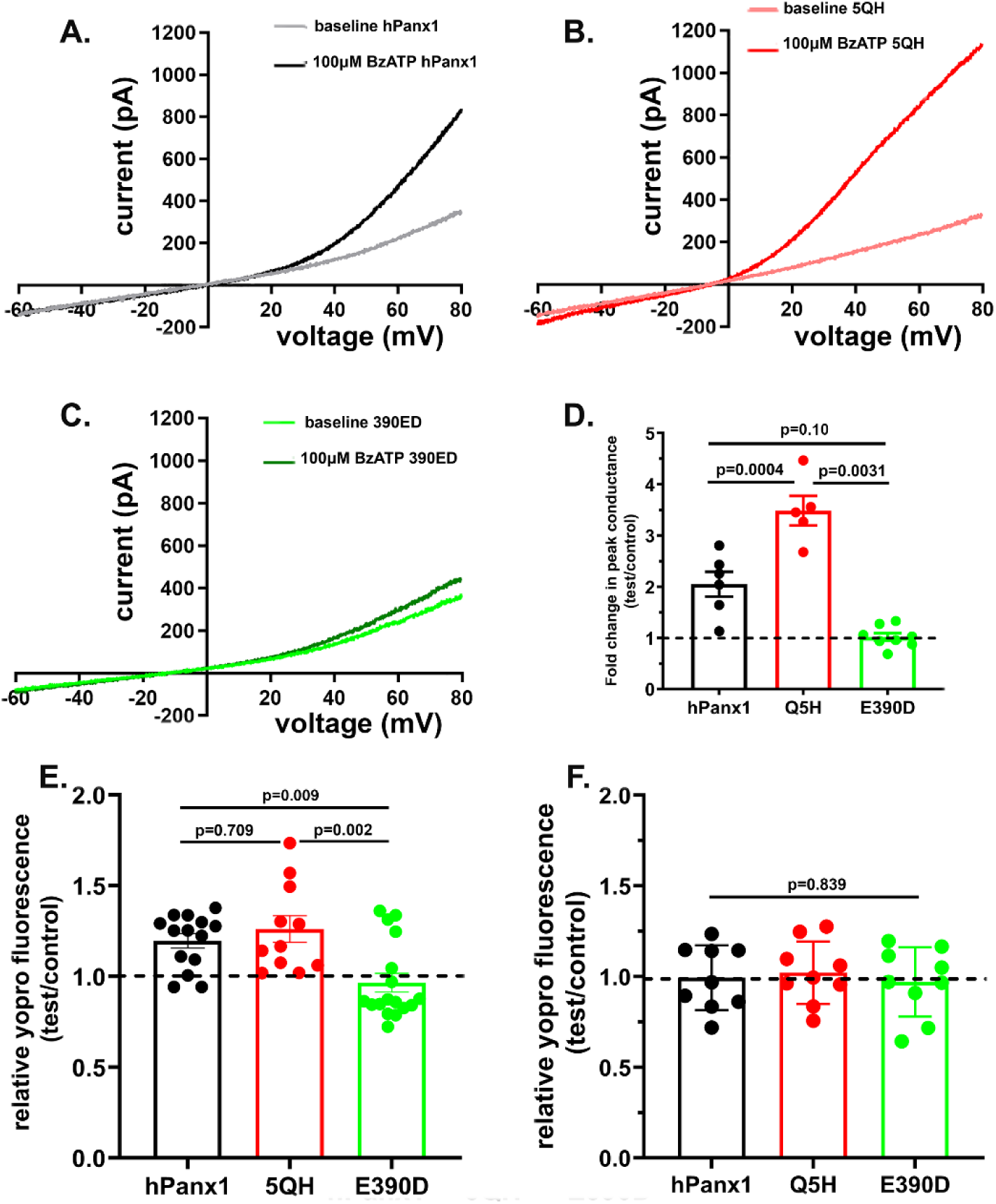
Differential activation of hPanx1 variants following P2X7R stimulation. (**A**-**C**) Examples of Panx1 currents recorded from N2a cells expressing hPanx1 (**A**) and two of its variants (Q5H: **B** and E390D: **C**) before and after 100 µM BzATP bath application. Note that N2a cells expressing hPanx1 and Q5H showed increased outward current following BzATP application while E390D did not. (**D**) Mean ± SEM values of fold changes in peak conductance obtained from hPanx1, Q5H and E390D Panx1 channels induced by 100 µM BzATP. N = 5 - 6 cells. Note the significantly higher peak conductance in Q5H compared to hPanx1 and the absence of conductance change in E390D expressing cells. (**E**-**F**) Mean ± SEM values of the relative YoPro fluorescence intensity changes obtained from N2a cells expressing the hPanx1 and two variants exposed for 10 min to 100 µM BzATP in the (**E**) absence and (**F**) presence of MFQ. Dashed line represents mean values of fluorescence recorded from cells exposed to control solution. Each point on the graphs represents a mean value of YoPro fluorescence recorded from all cells present in a field of view obtained from at 3 – 5 independent experiments. p values obtained from ANOVA followed by Tukey’s multiple comparison test.

Similarly to what we found in electrophysiological recordings, dye uptake assays revealed that application of BzATP (100 µM) to N2a expressing hPanx1 (see Supplementary Figure S1) and its variant Q5H induced influx of the dye YoPro which was absent in cells expressing the E390D variant (Figure 5E). Addition of P2X7 receptor agonist led to 1.20 ± 0.04 fold (n = 14) increase in YoPro fluorescence intensity in hPanx1 cells, to a 1.26 ± 0.07 fold (n = 11) increase in Q5H-Panx1 cells, and no significant change (0.97 ± 0.05 fold, n = 17) in E390D-Panx1 cells (Figure 5E). To further confirm the involvement of Panx1 in the uptake of the dye, we incubated the cells in the presence of the Panx1 channel blocker mefloquine (MFQ). As shown in Figure 5F, MFQ (10 µM) prevented the influx of YoPro induced by BzATP in hPanx1 (0.99 ± 0.06, n = 9) and in Q5H (1.02 ± 0.06, n = 9) and had no significant effect on E390D responses (0.97 ± 0.06, n = 9).

Evidence that the properties of hPanx1 channels exogenously expressed in N2a cells are similar to those found in native systems is provided in Supplemental Figure S2, which shows that the human astrocytoma cell line (1321N1, which lacks P2 receptors) are insensitive to K^+^ but respond to BzATP when expressing the rP2X7 receptor [36].

## DISCUSSION

Pannexins (Panx), which are the most newly recognized members of the gap junction family of proteins, include three members (Panx1-3). Panx1 has been shown to form channels in the cell membrane and shown to participate in physiological and pathological conditions distinct from those where connexin proteins are involved. Regarding its mode of activation, Panx1 channels have been shown to be gated by several stimuli, including membrane depolarization, membrane stretch and hypotonicity [3, 15, 16], elevated extracellular [K^+^] [9, 10, 37–39], and following the stimulation of many types of ligand-gated receptors (NMDA, P2X7, alpha-adrenergic, insulin, CXCR4) [2, 17–23]. Conductance and permeability states of Panx1 channels are reported to be stimulus dependent, such that activation by positive voltages leads to small conductance (∼80 pS) channels that are anion permeable mainly as Cl^-^ channels [39–41]. By contrast, mechanical stretch, elevated extracellular [K^+^], and P2 receptor stimulation cause reversible conformational changes in Panx1 channels leading to large conductance (∼500 pS) non-selective channel openings [42, 43]. However, differences between Panx1 orthologs have been reported, indicating that mPanx1 but not hPanx1 channels open upon membrane depolarization [24]. In addition, conflicting results exist with regards to the activation of Panx1 by high extracellular [K^+^], with studies showing that K^+^ sensitivity may be quite different among Panx1 orthologs and/or when studied in different expression systems (review in [25]).

Our present study performed on N2a cells expressing endogenous mPanx1 and on Panx1-null N2a cells expressing hPanx1 shows that hPanx1 channels generate outwardly rectifying currents when bathed in 2.5 mM K^+^ solution, although the mean peak conductance (∼2.0 nS) was significantly lower than that measured from cells expressing endogenous mPanx1 (∼4.0 nS). Activation of hPanx1 channels by membrane depolarization has been reported to occur only following caspase-3 cleavage of the carboxyl terminal domain at amino acid residue 378 [24]. However, this irreversible Panx1 activation after caspase cleavage does not lead to changes in single channel conductance or ion selectivity [24, 44]. Thus, considering that single channel conductance is reportedly similar for human and mouse Panx1 when activated by membrane depolarization, our data indicate that under our experimental conditions, fewer hPanx1 channels are open than mPanx1 channels.

Differences between hPanx1 and mPanx1 channel activation are not restricted to membrane depolarization but also with regard to K^+^ sensitivity. We here show that exposure to 10 mM K^+^ solution does not alter mean peak conductance in hPanx1 transfectants but doubled it in N2a cells expressing endogenous mPanx1. This difference between the two orthologs was also evidenced by the dye uptake method where influx of the YoPro dye was not induced in N2a expressing hPanx1 but was increased in N2a cells endogenously expressing mPanx1. Thus, these results suggest that the wide range of [K^+^] needed to activate Panx1 channels reported in the literature is most likely due to differences in K^+^ sensitivity between human and mouse Panx1. Indeed, our previous studies using primary cultures of mouse astrocytes and the human 1321N1 astrocytoma cell line showed that 10 mM K^+^ leads to dye influx through mPanx1 [9], while 50 mM K^+^ [37] but not 10 mM K^+^ (present study) was found to induce dye influx through the hPanx1 channels.

A recent study performed on oocytes and HEK293T cells exogenously expressing rat and mouse Panx1 reported a lack of activation of ethidium bromide uptake and outward current by 50 mM KCl [45]. That study also concluded that Panx1 channels possessed distinct gating mechanisms for ions and fluorescent dyes based on the observation that Panx1 channel blockers prevented or even enhanced Panx1 currents . Whereas a switch between permeability to anions vs cationic dyes has been proposed following Panx1 activation by modalities other than membrane depolarization (see [14, 39]), we found very similar fold changes in mPanx1 conductance and dye influx following channel activation by either high [K+] or BzATP, all of which were blocked by a pharmacological inhibitor. And we recently reported that cholesterol depletion enhances dye uptake and membrane currents in astrocyte and Panx1 transfectants, indicating similar effects of another gating stimulus [28]. Differences among studies of Panx1 activation, permeability and inhibition may have arisen from expression systems used, methods of analysis and fluorophores used and could be due to direct properties of the Panx1 pore or consequence of interaction with proteins bound to it.

Previous studies have proposed that the mechanism by which extracellular [K^+^] modulates Panx1 channel conductance and permeability is independent of membrane potential changes [9, 37] and instead is mediated by direct binding of this cation to the first extracellular loop of Panx1, likely at position R75 [46–48]. Given the virtually identical amino acid sequences in this region of the mouse and human Panx1 homologs, it seems unlikely that binding to this domain is the sole determinant of gating by extracellular K^+^. Further study is needed to fully define the domains differentially conferring K^+^ sensitivity to the homologs. Because truncated mPanx1 (mPanx1del378) displays similar conductance and anion permeability as full-length mouse and human Panx1 when activated by membrane depolarization, but remains sensitive to K^+^ [14, 44], it is likely that domains other than the carboxyl terminus may contribute to the gating process, either through inherent sequence sensitivity or mediated by possible binding partners.

Besides its activation by K^+^, rodent Panx1 channels are opened following stimulation of P2X7 [17, 18] and NMDA receptors [19] via phosphorylation by src-tyrosine kinase [2, 49] at position Y308 [50]. Activation of mPanx1 via receptor stimulation or extracellular K^+^ elevation leads to large pore formation and permeability to both anions and cations (reviewed in [14]). Here we show that mouse and human Panx1 are similarly sensitive to a P2X7 receptor agonist, BzATP, which increased mean peak conductance by about two-fold and led to 20-50% increase in the influx of YoPro. Interestingly, the two hPanx1 variants responded differently to BzATP stimulation. We found that the Q5H variant which has been previously reported to form hyperactive channels [26, 34, 35] displayed significantly higher mean peak conductance than that recorded from hPanx1 when stimulated with the P2X7 receptor agonist. On the other hand, however, because the E390D variant was totally insensitive to this mode of stimulation, it is likely that the channels this variant forms are functionally dead, possibly like those of rs143240087 (R217H) which was reported to have dominant-negative effects [27].

In conclusion, we show here that mouse and human Panx1 show different degrees of sensitivity to two well-known stimuli. These differences may have translational implications especially with regard to studies involving neurological disorders such as epilepsy. Given that extracellular [K^+^] increases to about 10-20 mM during seizures or spreading depolarization [51–53], the reported activation of hPanx1 channels during epileptiform activity [54] is in contrast to mPanx1 [10], likely due to stimuli other than K^+^, possibly involving a receptor mediated mechanism, such as by the NMDA receptor [19]. In addition, our characterization of the two most common *PANX1* variants in the human population suggests that differences in sensitivity/susceptibility to certain diseases may be related to the striking differences here described between the most common variant (Q5H) which is present in 71% of the population and the E390D which is expressed in 4% of the population.

## Acknowledgements

We thank Dr. Winfried Edelmann and Dr.Yongwei Zhang of the Einstein Gene Targeting and Core for providing CRISPR deletion of Panx1 in N2a cells and Dr. Randy Stout (currently at NYIT, Old Westbury NY) for his considerable assistance in design of human Panx1 constructs. We are grateful to Mr. Jian Pan (NYMC) for his technical assistance with cell culture and maintenance of frozen stocks. We acknowledge the contribution of Miss Xiaojun (Nancy) Feng who participated in preliminary experiments during the 2018 Summer Research Program at NYMC.

## Declaration of Interest

None

## Abbreviations

Panx1: Pannexin1
hPanx1: human Pannexin1
mPanx1: mouse Pannexin1
K^+^: potassium ion
N2a: neuroblastoma cells
1321N1: human astrocytoma cell line
FBS: Fetal Bovine Serum
CRISPR: Clustered Regularly Interspaced Short Palindromic Repeats
qPCR: quantitative polymerase chain reaction
BzATP: 3’-O-(4- Benzoyl)-benzoyl ATP
YoPro-1: Oxazole Yellow.

## Supporting information

Supplemental Figure S1. Representative images of dye uptake by N2a cells expressing endogenous mPanx1, lacking Panx1 (KO), and expressing hPanx1 after for 10 min exposure to saline (control) and to saline containing 10 mM K^+^ or 100 µM BzATP. Images were acquired from 4% PFA fixed cells. Scale bar: 20 µm.

Supplemental Figure S2. Endogenous hPanx1 channels are insensitive to elevated extracellular K^+^ but sensitive to P2X7 receptor activation. (**A**-**B**) Examples of hPanx1 currents recorded from 1321N1 human astrocytoma cell line expressing rP2X7 receptor exposed to solutions containing 2.5 and 10 mM K^+^ and to 100 µM BzATP. (**C**) Fold changes in peak conductance measured from 1321N1 cells exposed to high potassium (n=6) and BzATP (n = 5). Note that elevated extracellular K^+^ concentration did not lead to increase in Panx1 currents above baseline, while exposure to BzATP caused a 2 fold increase in peak conductance. (**D**-**E**) Examples and quantitation of hPanx1 currents recorded from 1321N1 cells exposed to high K^+^ and BzATP in the presence of 100 µM carbenoxolone (CBX, a gap junction and Panx1 channel blocker). Note in E that CBX prevented the increase in hPanx1 currents induced by BzATP. The generation and maintenance of 1321N1 cells expressing the rP2X7 were previously described in [36].

## Notes

### Competing Interest Statement

The authors have declared no competing interest.

